# HLA-Based Banking of Human Induced Pluripotent Stem Cells in Saudi Arabia

**DOI:** 10.1101/2023.09.16.557826

**Authors:** Maryam Alowaysi, Robert Lehmann, Mohammad Al-Shehri, Moayad Baadheim, Hajar Alzahrani, Doaa Aboalola, Asima Zia, Dalal Malibari, Mustafa Daghestani, Khaled Alghamdi, Ali Haneef, Dunia Jawdat, Fahad Hakami, David Gomez-Cabrero, Jesper Tegner, Khaled Alsayegh

## Abstract

Human iPSCs’ derivation and use in clinical studies are transforming medicine. Yet, there is a high cost and long waiting time for autologous iPS-based cellular therapy, and the genetic engineering of hypo-immunogenic iPS cell lines is hampered with numerous hurdles. Therefore, it is increasingly interesting to create cell stocks based on HLA haplotype distribution in a given population. In this study, we assessed the potential of HLA-based iPS banking for the Saudi population. First, we analyzed the HLA database of the Saudi Stem Cell Donor Registry (SSCDR), which contains high-resolution HLA genotype data of 64,315 registered Saudi donors at the time of analysis. We found that only 13 iPS lines would be required to cover 30% of the Saudi population, 39 iPS lines would offer 50% coverage and 596 for more than 90% coverage.

Next, As a proof-of-concept, we launched the first HLA-based banking of iPSCs in Saudi Arabia. Using clinically relevant methods, we generated the first iPSC line from a homozygous donor for the most common HLA haplotype in Saudi. The two generated clones expressed pluripotency markers, could be differentiated into all three germ layers, beating cardiomyocytes and neuronal progenitors. To ensure that our reprogramming method generates genetically stable iPSCs, we assessed the mutational burden in the generated clones and the original blood sample from which the iPSCs were derived using whole-genome sequencing. All detected variants were found in the original donor sample and were classified as benign according to current guidelines of the American College of Medical Genetics and Genomics (ACMG).

This study sets a road map for introducing iPS-based cell therapy in the Kingdom of Saudi Arabia.

## Introduction

Induced pluripotent stem cells (iPSCs) are a type of stem cell that can be generated from adult somatic cells by reprogramming them to a pluripotent state (Takahashi et al., 2007). Human iPSCs can indefinitely proliferate in the lab and be directed to differentiate into derivatives of all three germ layers (Takahashi et al., 2007; Yu et al., 2007). These two characteristics make iPSCs an attractive source of cells for cell therapy (Singh et al., 2015; Mandai et al., 2017). Upon their discovery, iPSCs were hailed as a promising alternative to human embryonic stem cells (hESCs), as they overcome the ethical problems associated with hESCs derivation and alleviate the risk of immunological rejection (Takahashi et al., 2007; Taylor et al., 2012). However, it became evident that developing autologous iPS-based cell therapy products for every patient is a laborious process that is currently prohibitively expensive and time-consuming (Habka, Mann et al. 2015; Huang, Liu et al., 2019; Bravery et al., 2015).

Alternatively, Human Leukocyte Antigen (HLA)-based banking of iPSCs for allogeneic cell therapy became a more attractive option (Taylor et al., 2005). HLA-matched cell therapy has been widely employed for hematopoietic stem cell transplantation for patients with blood cancers and other hematological disorders (Gragert et al., 2014; Park et al., 2012). However, HLA loci are highly polymorphic; therefore, generating thousands of iPS lines would be impractical. To mitigate this, it has been previously proposed that the generation of iPS cell stocks from carefully selected donors who are homozygous for the most common HLA haplotypes found in a given population, could offer coverage for every patient in need and could allow for the development of off-the-shelf cell therapy products (Taylor et al., 1993; Opelz and Dohler, 2007; Johnson et al., 2010; Opelz and Dohler 2010; Lee et al., 2018; Álvarez-Palomo, et al., 2021).

To evaluate the feasibility of HLA-based banking of iPSCs in Saudi Arabia, we analyzed the database of the Saudi Stem Cell Donor Registry (SSCDR), which is a registry established to facilitate patient-donor matching for hematopoietic stem cell transplantation. The SSCDR database contained 64,315 high-resolution HLA genotypes of registered Saudi citizens at the time of our analysis. We found that HLA-based banking of iPSCs may be a suitable strategy for pilot implementation and introduction of iPS-based cell therapy in Saudi Arabia. Additionally, we herein describe the establishment of the first two iPS lines from a Saudi donor who is homozygous for the HLA haplotype with the highest frequency in the population and provide maximal coverage. We describe the donor recruitment process, the reprogramming method to be used, and quality control tests that will be employed in the establishment of the HLA haplobank of iPSCs in Saudi Arabia.

## Results

### Identification of HLA Homozygous Donors in the Saudi Population

To identify potential homozygous donors, we examined the SSCDR HLA database for haplotype frequencies for HLA-A, HLA-B, and HLA-DRB1. Matching for these loci reduces the allograft rejection and diminishes the use of immunosuppressive drugs (Opelz and Dohler 2010). Our analysis showed that generating iPS lines from homozygous donors for the ten most frequent haplotypes can be expected to offer haplotype matching for 12.94% of the Saudi population (Table 1). We also performed a 5-locus based analysis of the SSCDR database and compared our results with those described by Alfraih et al., 2021,which yielded a good correspondence (Supplementary figure 1)(Table 1).

**Table 1:**
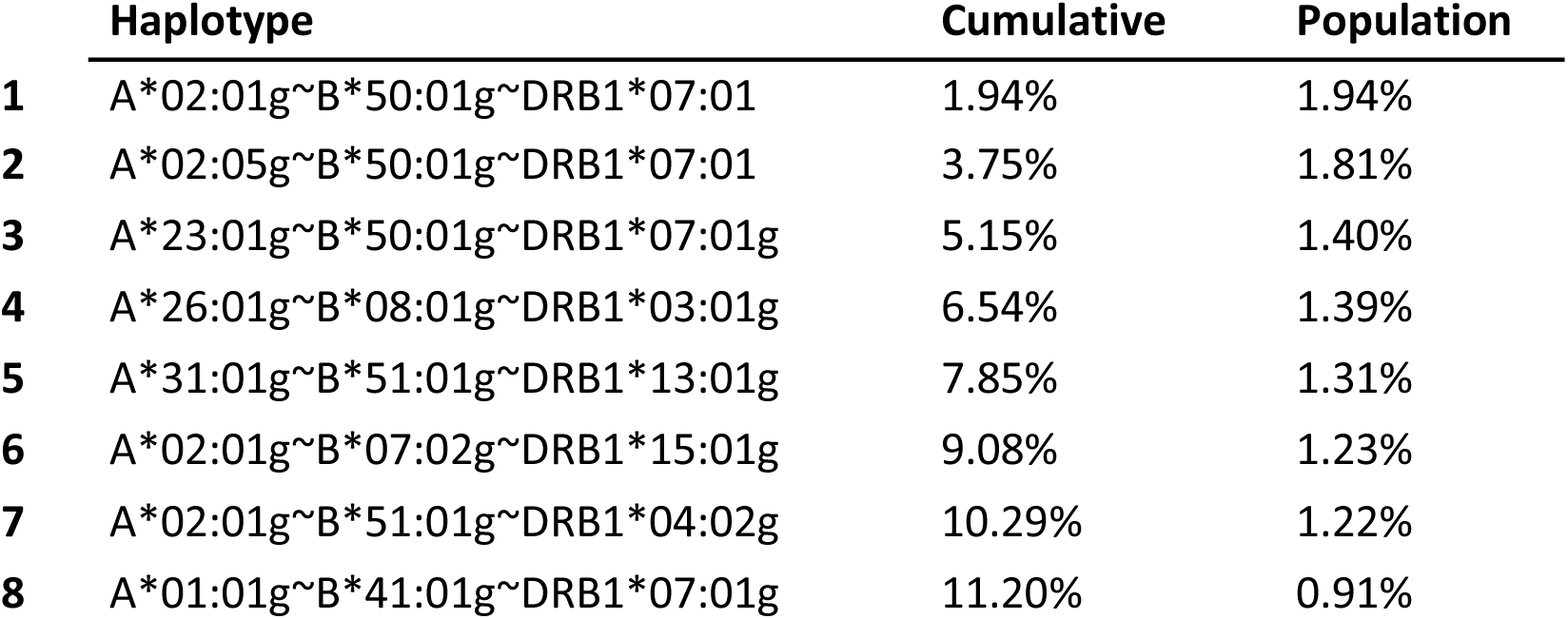
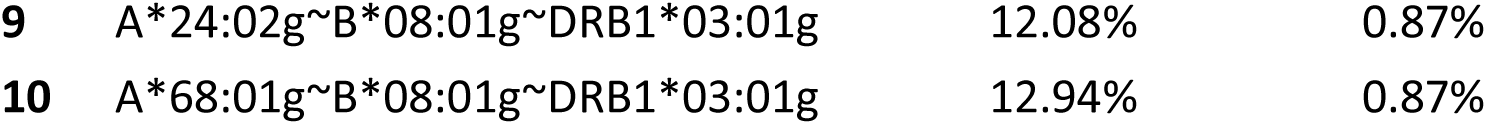
Frequencies and cumulative frequencies of the 10 most common HLA-A, HLA-B, and HLA-DRB1 in the Saudi Arabian population.

|  | Haplotype | Cumulative | Population |
| --- | --- | --- | --- |
| 1 | A*02:01g~B*50:01g~DRB1*07:01 | 1.94% | 1.94% |
| 2 | A*02:05g~B*50:01g~DRB1*07:01 | 3.75% | 1.81% |
| 3 | A*23:01g~B*50:01g~DRB1*07:01g | 5.15% | 1.40% |
| 4 | A*26:01g~B*08:01g~DRB1*03:01g | 6.54% | 1.39% |
| 5 | A*31:01g~B*51:01g~DRB1*13:01g | 7.85% | 1.31% |
| 6 | A*02:01g~B*07:02g~DRB1*15:01g | 9.08% | 1.23% |
| 7 | A*02:01g~B*51:01g~DRB1*04:02g | 10.29% | 1.22% |
| 8 | A*01:01g~B*41:01g~DRB1*07:01g | 11.20% | 0.91% |
| 9 | A*24:02g~B*08:01g~DRB1*03:01g | 12.08% | 0.87% |
| 10 | A*68:01g~B*08:01g~DRB1*03:01g | 12.94% | 0.87% |

Linkage disequilibrium (LD) scores between alleles of some of the most frequent haplotype in the Saudi population were shown to be low in some cases, suggesting a considerable possibility of recombination (Jawdat et al., 2020; Alfraih et al., 2021). We therefore modified the matching procedure by splitting haplotypes into individual loci and performing matching per locus, where each of the two possible alleles can be counted as potential match. Iterative selection and removal of the haplotype matching the most individuals (see Materials and Methods for details) yields a very similar order of the top ten haplotypes (Table 2), including only one new haplotype A*02:01g∼B*51:01g∼DRB1*03:01g. The fraction of HLA matches offered by these ten haplotypes, however, increases significantly to 26.9% (Table 2).

**Table 2:** Top 10 haplotypes that maximize the coverage across the population, using allele-wise matching across both haplotypes.

|  | Haplotype | Cumulative Coverage | Coverage | Number of Homozygous Donors |
| --- | --- | --- | --- | --- |
| 1 | A*02:01g~B*50:01g~DRB1*07:01 | 6.10% | 6.08% | 72 |
| 2 | A*02:05g~B*50:01g~DRB1*07:01 | 9.40% | 3.72% | 64 |
| 3 | A*26:01g~B*08:01g~DRB1*03:01 | 12.30% | 3.05% | 39 |
| 4 | A*31:01g~B*51:01g~DRB1*13:01 | 14.90% | 2.77% | 57 |
| 5 | A*02:01g~B*07:02g~DRB1*15:01 | 17.30% | 2.81% | 27 |
| 6 | A*02:01g~B*51:01g~DRB1*04:02g | 19.60% | 2.71% | 44 |
| 7 | A*23:01g~B*50:01g~DRB1*07:01g | 21.80% | 3.24% | 56 |
| 8 | A*02:01g~B*51:01g~DRB1*03:01g | 23.70% | 2.51% | 8 |
| 9 | A*24:02g~B*08:01g~DRB1*03:01g | 25.40% | 2.24% | 21 |
| 10 | A*01:01g~B*41:01g~DRB1*07:01g | 26.90% | 1.90% | 35 |

In extension, when using the maximized coverage approach, we found that a total of 13 haplotypes are estimated to have a match of 30% of the Saudi population (Figure 1), versus 51 lines, when selecting by maximum population frequency and using haplotype-wise matching. The number of required haplotypes covering 50% of the population increases to 39 and 220 for the maximum-coverage allele-wise, and maximum-frequency haplotype-wise approach, respectively. Since the generation of 39 iPS lines to cover >50% of the Saudi population is feasible, HLA-based banking of iPSCs may be a suitable strategy for the pilot implementation and introduction of iPS-based cell therapy in Saudi Arabia.

**Figure 1:**
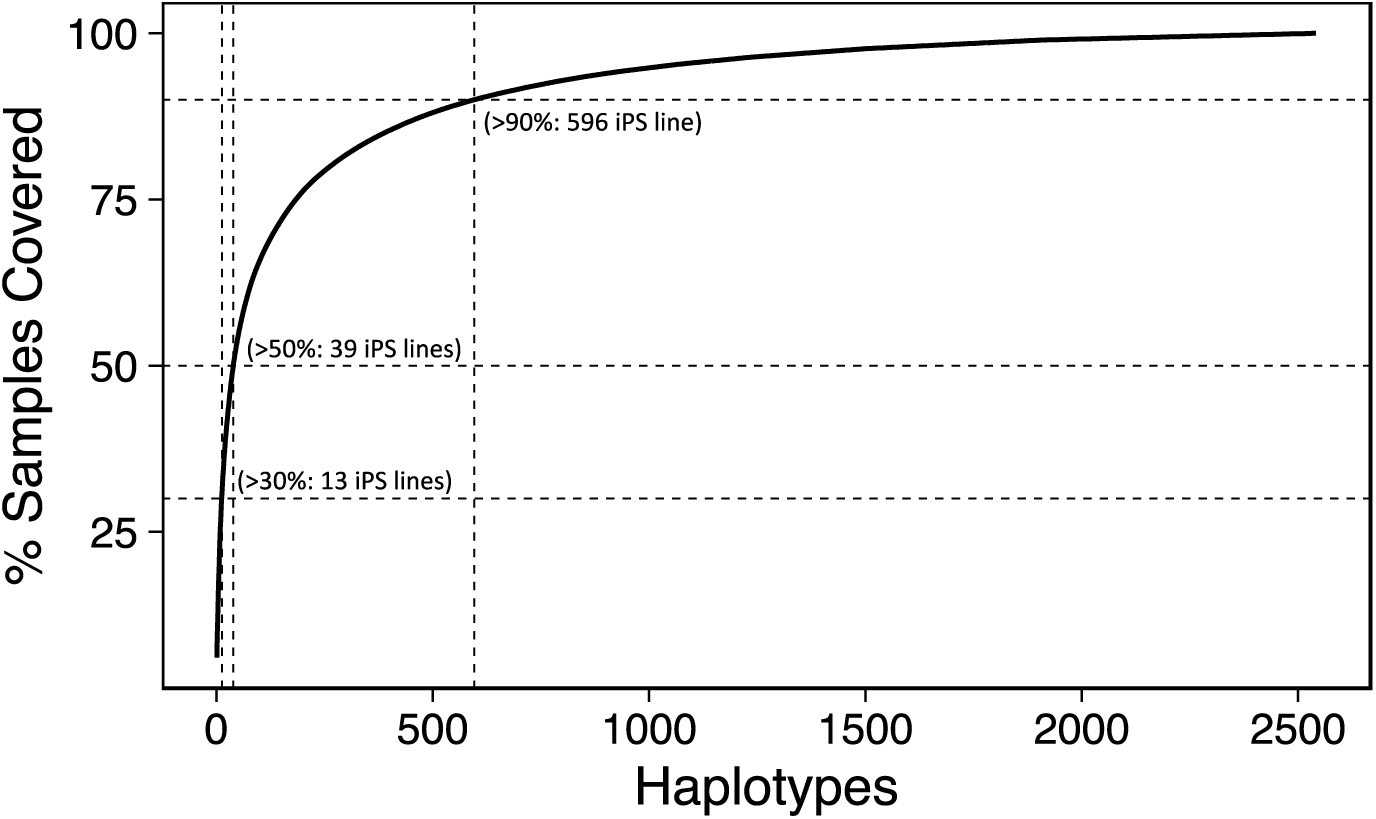
Estimated numbers of iPSC lines homozygous for HLA-A, HLA-B, and HLA-DRB1 (haplolines) and their coverage percentages for the Saudi population. The dotted lines mark 30, 50, and 90% coverage, for 13, 39 and 596 iPS lines, respectively.

### Donor recruitment and Derivation of HLA-haplobank iPS lines

In collaboration with the SSCDR, we identified a registered donor who is homozygous for the most common HLA haplotype (Table 1). This donor’s iPSCs would offer 6.1% coverage. The donor was initially contacted through the phone and upon approval, in-person interview was scheduled. After signing the informed consent, 10 ml peripheral blood sample was collected from the donor and erythroid progenitor cells (EPCs) were isolated expanded in culture for eight days. EPCs were chosen as the starting cell population for reprogramming due to their lack of DNA alterations and genomic structural variation including the absence of TCR/BCR genes recombination found in T-cells (Chou et al., 2011; Perriot et al., 2018; Araki et al., 2020).

Flow cytometry analysis showed that >69% of cells stained positive for the erythroid markers CD71 and CD235a after EPCs expansion (Figure 2A, 2B). On day 8 of expansion, reprogramming was initiated by electroporating EPCs with non-integrating episomal plasmids encoding OCT4, SOX2, KLF4, L-MYC, LIN28A, dominant-negative form of TP53, and EBNA1. At day 25 post transfection, numerous embryonic stem cell (ESC)-like colonies were identified with typical ESC morphological characteristics (i.e., distinct borders, bright centers, tight-packed cells, and a high nucleus-to-cytoplasm ratio) (Figure 2C). Such colonies were manually picked, expanded, and cryopreserved. Based on their ideal ESC-like morphology, two clones were chosen to be passaged and subjected to downstream pluripotency validation. The derived lines were registered in the Human Pluripotent Stem Cell Registry (hPSCreg) (https://hpscreg.eu/user/cellline/edit/KAIMRCi002-A, https://hpscreg.eu/user/cellline/edit/KAIMRCi002-B)

**Figure 2:**
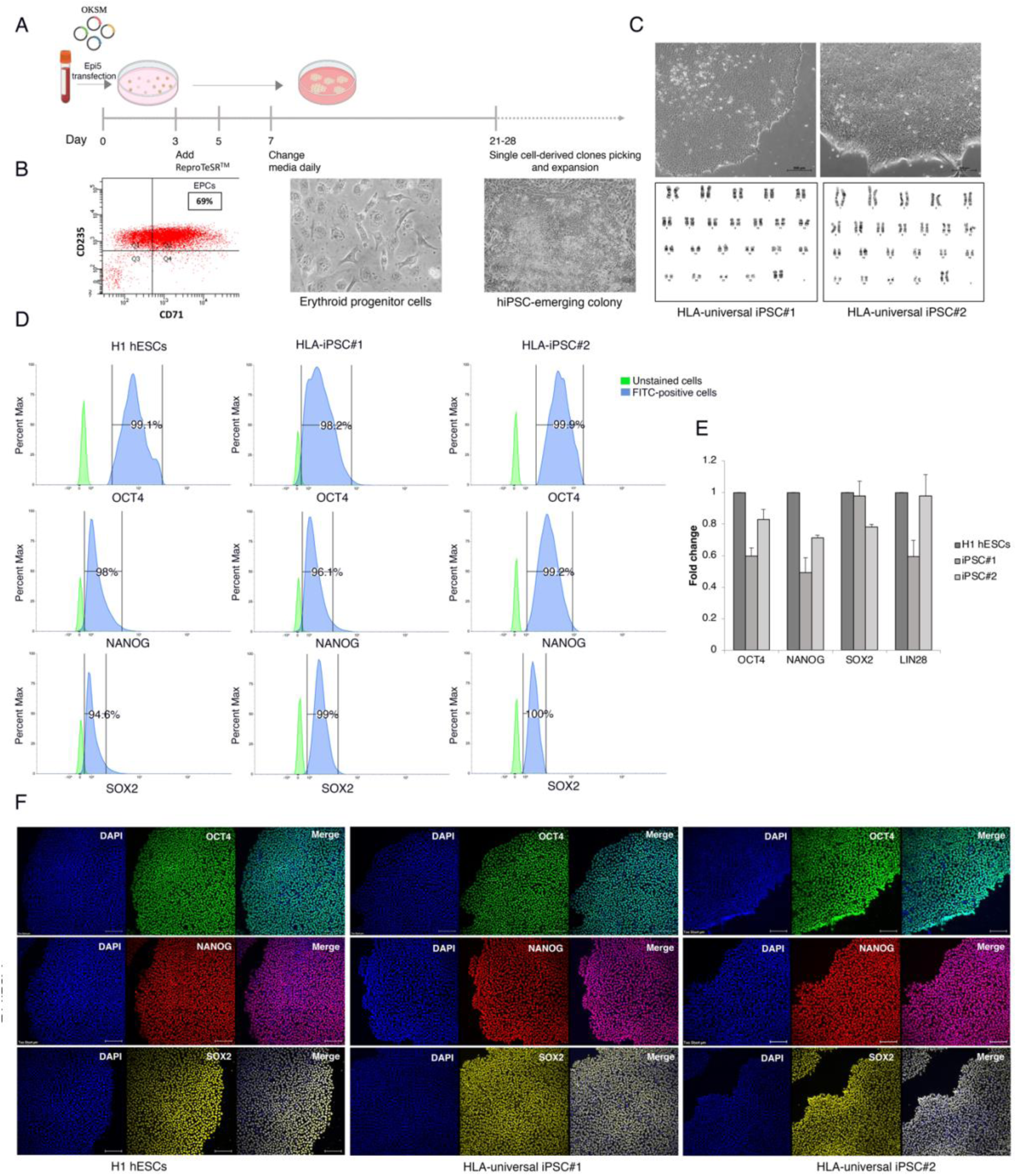
Generation and characterization of HLA-universal iPSCs. (A) Schematic representation of ReproTeSR^TM^ and episomal reprogramming method. (B) Flow cytometry histogram for erythroid markers CD71 and CD235a EPCs culture on day 8 shows that ∼70% of cells express the erythroid cell markers. Phase-contrast images of mesenchymal-to-epithelial transition and colonies appearance during reprogramming (days 11 to 28). (C) Top: representative images of HLA-universal iPS cell colonies generated from Erythroid Progenitor Cells exhibit more defined borders and compact morphology. Bottom: representative G-banded karyotype analysis for HLA-universal iPSCs shows normal karyotypes 46, XX. (D) Flow cytometry histograms of OCT4, NANOG, and SOX2 in HLA-universal iPSCs and H1 hESCs positive control. (E) Graph showing mRNA expression levels of pluripotency markers for the indicated iPSC lines presented as fold change relative to H1 hESC. Bars are median ± std of 3 biological replicates for each sample. (F) immunofluorescence staining of the pluripotency markers OCT4 (green), NANOG (red), and SOX2 (yellow), Nuclei were stained with DAPI (blue). Scale bar = 200 μm.

To assess the genomic integrity of HLA-iPSC#1 and #2, high-resolution G banding was performed after 12 passages in culture. More than 25 prometaphase spreads per clone were analyzed and showed normal female chromosomal number and structure (Figure 2C). Short tandem repeats (STR) assay confirmed the matching identity of the isolated iPS lines and the donor EPCs (Supplementary Figure 2A). Moreover, PCR analysis showed that the episomal plasmids were undetected in HLA-iPSC#1&2 after 12 passages (Supplementary Figure 2B). Additionally, mycoplasma testing showed that the generated iPSC lines are mycoplasma-free (Supplementary Figure 2C).

### Validation of iPSCs’ self-renewal and pluripotency properties

Pluripotency markers OCT4, NANOG, and SOX2 were detected at the mRNA and protein levels in both clones. Flow cytometry histograms demonstrated that >98% of cells stained positively for OCT4, >96% for NANOG, and >94% for SOX2 (Figure 2D). Moreover, the derived iPSC lines displayed positive expression of *OCT4, NANOG, SOX2*, and *LIN28* by RT-qPCR (Figure 2E) and OCT4, NANOG, and SOX2 by immunofluorescence (Figure 2F). Direct in-vitro differentiation to the three germ layers, mesoderm, definitive endoderm, and ectoderm was used to demonstrate the tri-lineage differentiation capacity. We observed a down-regulation of *OCT4* and *NANOG* and an upregulation of germ layer-specific markers by RT-qPCR (Figure 3B). Immunostainings for the neural progenitor marker (NESTIN) indicated ectodermal differentiation. The positive expression of Brachyury, a member of the Tbox family, showed an early determination of mesoderm. We further assessed the presence of the endodermal marker SRY-Box Transcription Factor 17 (SOX17) (Figure 3A). We, therefore, proved that the constructed HLA-universal iPSC lines possess bona fide characteristics of pluripotent stem cells. All performed quality control tests are summarized in Table 3.

**Figure 3:**
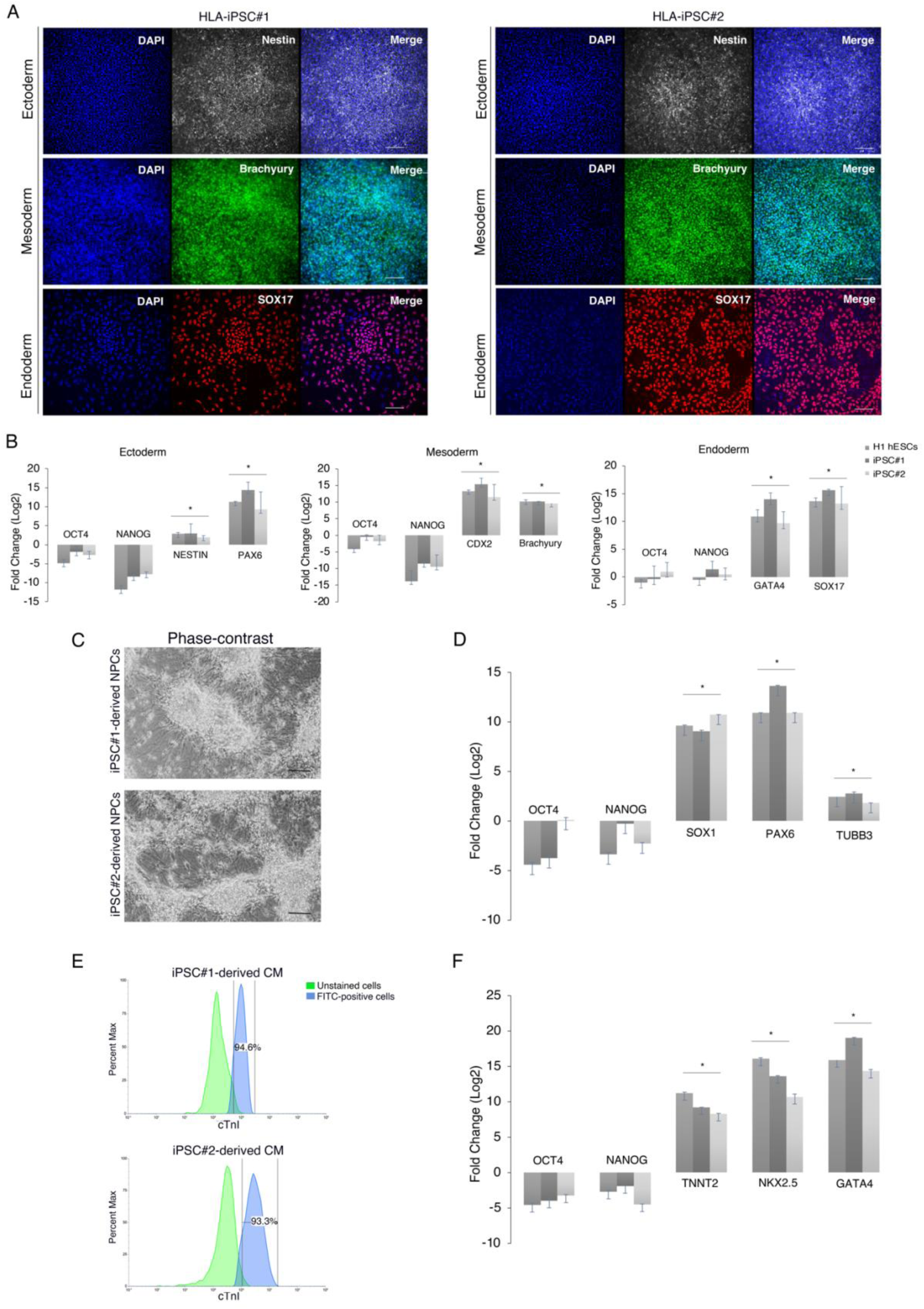
Differentiation potential of HLA-universal iPSCs. (A) immunofluorescence staining of specific markers for the three germ layers Ectoderm (NESTIN), Mesoderm (Brachyury), Endoderm (SOX17), Nuclei were stained with DAPI (blue). Scale bar = 200 μm. (B) Graphes showing mRNA expression levels of the lineage-specific markers for the three germ layers Mesoderm (*CDX2* and *Brachyury*), Endoderm (*GATA4* and *SOX17*), and Ectoderm (*NESTIN* and *PAX6*) presented as fold change relative to undifferentiated cells. Bars are median ± std of 3 biological replicates for each sample. Student’s t-tests, *p < 0.05. (C) Phase-contrast images of CNS-type NPC differentiation display typical NPC morphology 34 days post differentiation. Scale bar = 200 μm. (D) Graph showing mRNA high expression levels of CNS-type NPCs markers SOX1, PAX6, and a low or negative expression of β-tubulin III presented as fold change relative to undifferentiated cells. Bars are median ± std of 2 biological replicates for each sample. Student’s t-tests, *p < 0.05. (E) Flow cytometry histograms display positive cTnI expression in iPSC-derived CM. (F) Graphes showing mRNA expression levels of cardiac markers *TNNT2*, *NKX2.4*, and *GATA4*. Bars are median ± std of 2 biological replicates for each sample. Student’s t-tests, *p < 0.05.

**Table 3:** Summary of characterization tests performed on HLA-iPSC lines#1 and #2.

| Classification | Test | Results | Data |
| --- | --- | --- | --- |
| **Morphology** | Phase-contrast imaging | Typical primed pluripotent human stem cell morphology | Figure 2C, Top |
| **Karyotype** | G-banding | 46, XX | Figure 2C, Bottom |
| **Pluripotency status** | Quantitative analysis (Flow cytometry, RT-qPCR) | Flow Cytometry: 98% OCT4, 96% NANOG, 99% SOX2.<br>RT-qPCR: positive expression for OCT4, NANOG, SOX2, LIN28. | Figure 2D, 2E |
|  | Qualitative analysis (Immunocytochemistry) | Positive for the pluripotency markers: OCT4, NANOG, and SOX2. | Figure 2F |
| **Genetic identity** | STR profiling | 24 loci tested, all matched between donor EPCs and derived iPSC lines | Supplementary Figure 2A |
| **Verification of the absence of episomal vectors** | PCR analysis | The five episomal DNA plasmids were undetected after 12 passages | Supplementary Figure 2B |
| **Mycoplasma testing** | RT-qPCR | Negative | Supplementary Figure 2C |
| **Multilineage differentiation potential** | Directed differentiation | RT-qPCR measurement of expression levels of ectoderm (*PAX6* and *NESTIN*), mesoderm (*BRACHYURY* and *CDX2*), and endoderm (*SOX17* and *GATA4*). | Figure 3A, 3B |
|  |  | Immunocytochemistry: Positive for<br>NESTIN, BRACHYURY, and SOX17. |  |
| **HLA typing** | WGS | Homozygous at class I loci A, B, C, and class II loci DQB1 and DRB1, with only DPB1 being heterozygous | Figure 1 |
| **SV analysis** | WGS;<br>Manta+Delly+SURVIVOR | No SVs in iPSC#2; one tandem duplication in iPSC#1 | Supplementary<br>Figure 3 |
| **SNP analysis** | WGS; GATK, Mutect2 | No high-impact SNP acquired during cell line establishment; SNPs in genetic background of the donor classified as benign | Supplementary<br>table 3.<br>Table 4. |

Furthermore, the differentiation potential of the iPSC lines toward central nervous system (CNS)-type neural progenitor cells (NPCs) and beating cardiomyocyte was tested. CNS-type NPC differentiation induced a marked increase in key neuronal markers such as *SOX1*, *PAX6* and *TUBB3* (Figure 3C, 3D).

The HLA-iPSC clones were also differentiated into cardiomyocytes (CMs) through a step-wise protocol. By day 15, the iPSC-derived CMs displayed spontaneous contractions, a unique functional property of pulsating heart muscles (Supplemental file# movie). Flow cytometry histograms showed that >94% of cells stained positively for Cardiac Troponin I (cTnI) (Figure 3E). Moreover, RT-qPCR demonstrated that iPSC-derived CMs expressed the Cardiac Muscle Troponin T (*TNNT2*), myocardial markers NK2 Homeobox 5 (*NKX2.5*), and GATA family of zinc-finger transcription factor (*GATA4*) at mRNA levels (Figure 3F).

### Whole-Genome Sequencing of Generated iPS Lines

To ascertain the genotype and whether the genomic integrity of the constructed iPSC lines was maintained during reprogramming and prolonged cultivation, we sequenced the genomes of the parental blood sample and the progenies iPSC#1 and iPSC#2 at passage 13. Genotyping of the HLA loci using the 23x-26x coverage read datasets confirmed homozygous status at class I loci A, B, C, and class II loci DQB1 and DRB1, with only DPB1 being heterozygous (A*02:01∼A*02:01∼B*50:01∼B*50:01∼C*06:02∼C*06:02∼DPB1*02:01∼DPB1*04:01∼DQB1*02:0 2∼DQB1*02:02∼DRB1*07:01∼DRB1*07:01). Genomic variants were called in parental and progeny samples using GATK. This yielded a total of 5.4 million polymorphic sites with a mean genotype call rate of 99.2% and a heterozygosity ratio of 1.7. Out of the 4.3 million single nucleotide polymorphisms each sample had about 3.9 million polymorphic loci (Supplementary Table 2). We first focused on SNPs that were found polymorphic in all three samples to generate a high confidence variant set for the genetic background of the donor. We then examined any variants that might affect cancer-related genes based on the COSMIC Cancer Gene Census database and Shibata list as described previously (Yoshida et al., 2022) which involved 15 heterozygous SNVs and 4 homozygous SNVs in (see Materials and Methods for details). However, in the categories of sequence variants developed by the American College of Medical Genetics and Genomics (ACMG), we found that the 19 variants are almost certainly benign (Table 4). Thus, the direct link between these mutations and tumorigenicity was eliminated since the HLA universal donor was healthy at the time of iPSC generation.

**Table 4:** Examined variant loci in donor genetic background. Variant locus provided as [chromosome]:[bp], RefSeq transcript, Ensembl gene identifier, dbSNP variant identifier, genotype in donor (heterozygous/homozygous) and classification as per ACMG.

| Variant Locus | RefSeq | Gene | dbSNP | Genotype | ACMG Classification |
| --- | --- | --- | --- | --- | --- |
| 1:150853154-150853154 | ARNT (NM_001668.4):c.138-348C>T (p.?), Chr1(GRCh37):g.150825630G>A | ENSG00000143437 | rs4072037 | Heterozygote | Benign |
| 1:155192276-155192276 | MUC1 (NM_002456.6):c.66G>T (p.Thr22=), Chr1(GRCh37):g.155162067C>A | ENSG00000185499 | rs4072037 | Homozygote | Benign |
| 12:6590204-6590204 | CHD4 (NM_001273.5):c.3340+1261dup (p.?), Chr12(GRCh37):g.6699378dup | ENSG00000111642 | rs58925722 | Heterozygote | Benign |
| 14:60658222-60658222 | SIX1 (NM_005982.4):c.-9033T>C (p.?), Chr14(GRCh37):g.61124940A>G | ENSG00000126778 | rs10143202 | Heterozygote | Benign |
| 15:88253479-88253479 | NTRK3-AS1 (NR_038229.1):n.749+1G>A, Chr15(GRCh37):g.88796710G>A | ENSG00000260305 | rs1105693 | Heterozygote | Benign |
| 17:7667260-7667260 | TP53 (NM_000546.6):c.*2348dup (p.?), Chr17(GRCh37):g.7570591dup | ENSG00000141510 | rs34103303 | Heterozygote | Benign |
| 2:29221229-29221229 | ALK (NM_004304.5):c.3516-394G>A (p.?), Chr2(GRCh37):g.29444095C>T | ENSG00000171094 | rs1569156 | Homozygote | Benign |
| 3:10046723-10046724 | FANCD2 (NM_001018115.3):c.1278+1del (p.?), Chr3(GRCh37):g.10088408del | ENSG00000144554 | rs750338758 | Heterozygote | Benign |
| 3:149657502-149657502 | WWTR1-AS1 (NR_040250.1):n.-708G>A, Chr3(GRCh37):g.149375289G>A | ENSG00000018408 | rs6783790 | Heterozygote | Benign |
| 5:142771377-142771377 | ARHGAP26 (NM_001135608.3):c.154+462T>C (p.?), Chr5(GRCh37):g.142150942T>C | ENSG00000145819 | rs10042074 | Heterozygote | Benign |
| 5:143214087-143214087 | ARHGAP26 (NM_001135608.3):c.2190C>T (p.Asn730=), Chr5(GRCh37):g.142593652C>T | ENSG00000145819 | rs258819 | Homozygote | Benign |
| 6:29943463-29943463 | HLA-A (NM_002116.8):c.539T>A (p.Leu180X), Chr6(GRCh37):g.29911240T>A | ENSG00000206503 | rs9260156 | Heterozygote | Benign |
| 6:29944251-29944252 | HLA-A (NM_002116.8):c.751del (p.Asp251fs), Chr6(GRCh37):g.29912030del | ENSG00000206503 | rs45576436 | Heterozygote | Benign |
| 6:41936060-41936060 | CCND3 (NM_001760.5):c.759G>T (p.Glu253Asp), Chr6(GRCh37):g.41903798C>A | ENSG00000112576 | rs33966734 | Heterozygote | Benign |
| 6:149716206-149716206 | LATS1 (NM_004690.4):c.-141+1643G>A (p.?), Chr6(GRCh37):g.150037342C>T | ENSG00000131023 | rs182844352 | Heterozygote | Benign |
| 7:116724769-116724770 | MET (NM_000245.4):c.1201-6898del (p.?), Chr7(GRCh37):g.116364824del | ENSG00000105976 | rs34822187 | Heterozygote | Benign |
| 7:152247986-152247986 | KMT2C (NM_170606.3):c.2447dup (p.Tyr816X), Chr7(GRCh37):g.151945072dup | ENSG00000055609 | rs150073007 | Heterozygote | Benign |
| 9:134029563-134029564 | BRD3OS (NM_001355256.2):c.*2564del (p.?), Chr9(GRCh37):g.136894688del | ENSG00000235106 | rs139424439 | Heterozygote | Benign |
| X:41357831-41357831 | DDX3X (NM_001356.5):c.*10112A>T (p.?), ChrX(GRCh37):g.41217084A>T | ENSG00000215301 | rs6520743 | Homozygote | Benign |

In the second step we tested whether the cell lines acquired new SNPs compared to the donor, using the donor sample as matched normal for the cell line samples. This approach yielded 1,610 and 1,888 SNPs for iPSC#1 and iPSC#2, respectively (Supplementary Table 3). None of the detected SNPs is predicted to have high impact with the majority classified as modifier.

While the subtractive analysis of structural variants (SVs) of donor vs. cell line did not detect newly acquired mutations in HLA-iPSC#2, it revealed a heterozygous tandem duplication on chromosome 16 (74,726,891bp – 74,727,373bp) in HLA-iPSC#1 (Supplementary Figure 3) which spans part of exon 3 of the Fatty Acid 2-Hydroxylase (FA2H) where it could lead to an alteration in the transcript. However, this gene is not part of the COSMIC Cancer Gene Census database and Shibata list rendering this variant benign.

## Discussion

Within only seven years of their initial derivation in 2007, iPSCs moved to clinical studies when a patient with age-related macular degeneration (AMD) was the first recipient of autologous iPS-derived retinal pigment epithelial cell sheet, in the world’s first in human clinical trial (Mandai et al., 2017). However, it became evident that the high cost and extended waiting time associated with autologous iPS-based cellular therapy, posed a significant hurdle to the advancement into the clinical domain (Habka, Mann et al. 2015; Huang, Liu et al., 2019; Bravery et al., 2015).

One approach that was proposed to solve the time and cost problems, is the creation of a hypo-immunogenic iPS cell line that evades the immune system. In this approach, iPS cells would be genetically modified to inactivate major histocompatibility complex (MHC) class I and II genes (Xu et al., 2019; Kitano et al., 2022). However, to achieve this, multiple rounds of gene editing are required, which extends the time the cells are cultured, thus increasing the risk of acquiring mutations. Additionally, gene editing technologies like, CRISPR/Cas9 has been shown to introduce unintended genomic aberrations and may render the cells not useful for therapy (Fu et al., 2013; Cradick et al., 2013). Even base and prime editing that does not involve double strand breaks have recently been shown to induce significant genotoxicity in human cells (Fiumara et al., 2023).

Therefore, there has been an increased interest in HLA-based iPS banking in numerous countries (Taylor et al., 2012; Gourraud et al., 2012; Lee et al., 2018; Álvarez-Palomo, et al., 2021; Yoshida et al., 2023). In this study, we assessed the feasibility of creating an iPS haplobank in Saudi Arabia to develop clinical-grade iPS cell stocks, as the ultimate goal. In order to achieve this, we used the high-resolution HLA genomic database of the SSCDR, which at the time of analysis contained 64,315 registered donors, and assumed it was a representative sample of the Saudi population. We found that, the feasibility of HLA-based banking in Saudi Arabia is comparable to similar endeavors in other countries. We found that an iPS haplobank of the top 5 haplotypes that offer maximal coverage for the Saudi population would cover 17.30% of the population, which is close to the Spanish bank in which, the top 5 haplotypes cover 19.44%, but lower than the Korean estimation, in which the top 5 haplotypes cover 27.99% (Lee et al., 2018; Álvarez-Palomo, et al., 2021). This finding is in line with previous reports that showed a relatively high HLA genetic diversity among Saudis compared to other populations (Alfraih et al., 2021).

We found that an iPS haplobank generated from homozygous donors from the top 39 haplotypes would offer coverage of more than 50% of the Saudi population. This significant percentage may allow for many Saudi patients to benefit from iPS-derived cell therapies in the kingdom and therefore, it justifies the construction of the haplobank. In addition, streamlining the process of generating clinical-grade iPSCs will facilitate the establishment and future expansion of the bank to include additional haplotypes. Moreover, due to high level of consanguinity in the Saudi population, there is a considerable excess homozygosity, which may facilitate the identification of homozygous donors and haplobanking (Chentoufi et al., 2022).

Due to the relatively high intra-population diversity in Saudi, we found that achieving higher coverage requires much larger cell stocks. Around 596 iPS line would be required to cover 90% of the population, and 2541 lines for 100% coverage. Even though we envisage that the establishing of an iPS cell stock to cover 30%-50% of the Saudi population is a feasible goal to introduce iPS-based cell therapy in the kingdom, achieving higher coverage percentage becomes increasingly cost-ineffective. Therefore, more research is needed to improve current methods of clinical-grade iPS generation to reduce cost and waiting time to make autologous cell therapy a possibility. Additionally, as we gain tighter control on the outcome of gene editing technologies, the creation of clinically-relevant universal hypoimmunogenic iPS lines might become more feasible in the future.

To establish the workflow and initiate HLA-based banking in Saudi, we recruited the first donor and generated the first two clinically relevant iPS lines using defined feeder-free conditions. We chose EPCs to be the starting cell population for reprogramming. As opposed to human dermal fibroblasts, EPCs can be easily isolated and expanded from a simple ten ml blood sample and does not require painful skin biopsies. This is of particular importance in the donor recruitment process, as participants might be discouraged to donate if the procedure is invasive.

Additionally, EPCs are frequently replenished in the blood and therefore are less likely to accumulate environment-induced mutations like fibroblasts (Panther et al., 2021; Kamimura et al., 2021). Moreover, they lack the TCR/BCR genes recombination found in T-cells, making them a more attractive source of iPS cells. This is in addition to recent research demonstrating that erythroblasts-derived iPS cells are less likely to harbor genetic aberrations when compared to iPS cells from other sources (Perriot et al., 2018; Araki et al., 2020).

Eight days of expansion showed that around 69% of the cells were CD71^+^CD235a^+^ (Figure 2B). The rest were CD71^-^CD235a^+^ and are more likely to be differentiated erythroblasts on their way to enucleation and are therefore, unamenable to reprogramming. Differentiated cells were particularly evident as red colored cells when EPCs were pelleted by centrifugation.

As an alternative to conventional retroviral-based cell reprogramming, non-viral, non-integrating plasmid-based reprogramming technique is more clinically-relevant (Yu et al. 2009; Okita et al., 2011; Bang et al., 2018). The reprogramming factors are delivered by vectors that contain oriP and EBNA-1, based on the Epstein-Barr Nuclear Antigen-1, which has demonstrated the ability to produce iPSCs highly efficiently without the potential risk of transgenic sequences being inserted into the target cell genome (Drozd et al., 2015). As opposed to other non-integrating reprogramming methods like sendai virus and mRNA, episomal plasmids is the most cost-effective. Additionally, we found that these plasmids are readily removed from the reprogrammed cells as they were expanded, with most lines testing negative by end-point PCR by passage 12.

Following the expansion of EPCs, electroporation of the reprogramming episomal was carried out. ESC-like colonies appeared around 20-25 days post transfection and were characterized by distinct borders, bright centers, tight-packed cells, and a high nucleus-to-cytoplasm ratio. The iPS clones were mechanically picked, expanded, and characterized for self-renewal and pluripotency in feeder-free culture conditions.

For developing clinical-grade HLA haplobank, KAIMRC is currently establishing its cell-processing-center in compliance with the good manufacturing practice (GMP) guidelines. Although the current HLA-iPSC#1 and iPSC#2 were generated inside research-grade labs, future haplobanking and clinical products will be derived and cryopreserved inside our GMP facility including re-derivation of the current HLA-iPSC lines to clinical-grade. Re-derivation of human pluripotent stem cell lines inside GMP facilities has been done before. For instance, the H1 hESC line was re-derived and used as part of Astellas Pharma’s phase II retinal pigment epithelium (RPE) trial, and re-derived H9 hESC line was used to generate dopaminergic neurons for a Parkinson’s disease clinical trial by BlueRock Therapeutics (Sullivan a et al., 2020).

## Conclusions

Our study lays the foundation for the roadmap towards HLA-based banking of human induced pluripotent stem cells (iPSCs) in Saudi Arabia (Figure 4). By interrogating the Saudi Stem Cell Donor Registry, we identified a subset of homozygous donors that could offer considerable coverage for the Saudi populace. Our analysis revealed that achieving 30% and 50% coverage necessitate the generation of 13 and 39 iPS lines, respectively. As a proof of principle, we successfully generated the first HLA-iPS line (2 clones), that offer 6.1% coverage of the Saudi population. By employing clinically-relevant methodologies, we ensured the safety and quality of these iPSCs. Notably, whole-genome sequencing confirmed the genomic stability of the generated lines, hence alleviating concerns of high-risk mutations during reprogramming and expansion processes. Our study substantiates the feasibility of HLA-based iPSC banking in Saudi Arabia and paves the way for a resilient infrastructure in regenerative medicine and personalized therapeutics.

**Figure 4:**
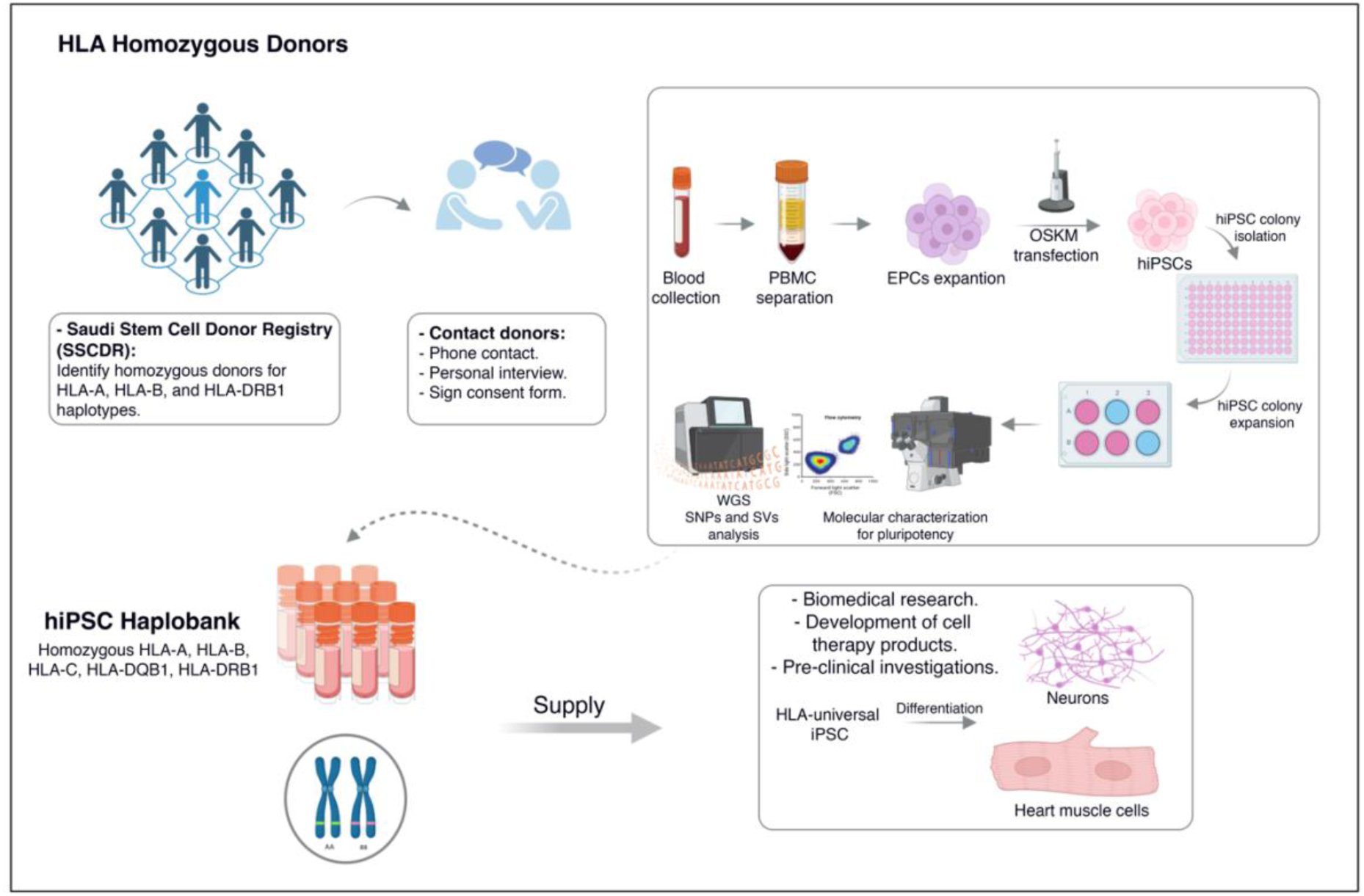
Graphical summary of the undergoing HLA-based banking in Saudi Arabia.

## Supplementary Information

Additional file 1: Supplementary Figure 1. Alfraih et al haplotype frequency.

Additional file 2: Supplementary Figure 2. HLA-iPSC lines authentication.

Additional file 2: Supplementary Figure 3. Tandem duplication on chromosome 16 in cell line iPSC#1 visualized using IGV.

Additional file 3: Supplementary table 1. Comparison of 5-locus based haplotype frequencies based on the SSCDR database and KFSH&RC dataset (Alfraih et al. 2021).

Additional file 4. Supplementary table 2. Single nucleotide polymorphisms for donor blood sample and cell lines iPSC#1 and iPSC#2. The fraction of genotype calls matching with donor was calculated for biallelic SNPs.

Additional file 6. Supplementary Table 3: Subtractive SNP calls for cell lines iPSC#1 and iPSC#2 vs. donor.

## List of abbreviations

SSCDR: Saudi Stem Cell Donor Registry
HLA: Human leukocyte antigen
EPCs: Erythroid progenitor cells (EPCs)
hESC: Human embryonic stem cell
iPSC: Induced pluripotent stem cells
hPSCreg: Human Pluripotent Stem Cell Registry
WGS: Whole Genome Sequencing
SNPs: Single nucleotide polymorphisms
SVs: Structural variants
AMD: Age-related macular degeneration
GMP: good manufacturing practice
RPE: Retinal pigmented epithelial cells

## Acknowledgements

We thank King Abdullah International Medical Research Center (KAIMRC) and the Saudi Stem Cell Donor Registry (SSCDR) for facilitating the initiation of HLA-based iPS banking. We thank current, and future donors for their valuable donations.

## Authors’ contributions

KA contributed through the conception of the idea, the design of the work, interpretation of data, and wrote the manuscript. MA contributed through sample processing, iPS generation, validation assays and differentiation and writing the manuscript. RL designed the iterative algorithm and analyzed the SSCDR HLA database to identify haplotypes and coverage, WGS data analysis and writing the manuscript. MA, MB, HA, DA, AZ, DM, have all contributed in iPS derivation and validation tests. MD performed karyotype analysis. Khaled Alghamdi performed the STR tests. DJ supervised the data curation of the SSCDR. FH assessed the clinical significance of identified variants revealed by WGS. DGC, JT and AH have participated in the design of the work and revision of the document.

## Funding

This work is funded by KAIMRC grant RJ20/134/J. Additional funding was provided through KAUST’s Smart Health Initiative.

## Availability of data and materials

All the presented data is available for consultation.

## Declarations

### Ethics approval and consent to participate

The interrogation of the HLA database of the SSCDR, informed consents and donor sample collection were approved by the institutional review board (IRB) and research ethics committee of KAIMRC (RJ20/134/J).

### Competing interests

The authors declare no competing interests.

### Figure legends

**Supplementary** Figure 1: Comparison of cumulative 5-locus haplotype frequency of the SSCDR HLA database and the Alfraih et al.

**Supplementary** Figure 2: Cell lines authentication. (A) Short Tandem Repeat (STR) profiling guaranteed the genetic identity between the established iPSC lines and the donor EPCs. (B) PCR analysis detected the absence of the episomal plasmids in the indicated lines at passage 12. (C) RT-qPCR showing negative mycoplasma test in HLA-iPSC lines.

**Supplementary** Figure 3: Tandem duplication on chromosome 16 in cell line iPSC#1 visualized using IGV.

## Materials and Methods

### Haplotype frequency analysis

HLA haplotype frequencies in the Saudi Arabian population were estimated based on haplotype information stored in the Saudi Stem Cell Donor Registry (https://kaimrc.ksau-hs.edu.sa/?page_id=1481) database. This database, comprising 64,315 individuals at the time of analysis, was analyzed using the EM algorithm as implemented in Hapl-o-Mat v 1.1 (DOI: 10.1007/978-1-4939-8546-3_19) to estimate population level haplotype frequencies using two digit resolution. The haplotype coverage was estimated as detailed in Álvarez-Palomo (https://doi.org/10.1186/s13287-021-02301-0) using an iterative algorithm. In each iteration, the most frequent haplotype was identified and all matching individuals were counted and removed from the dataset before the next iteration on the remaining dataset. Importantly, the haplotype matching procedure was modified by considering each locus independently and allowing matches on either of the two possible alleles per locus. Matching was performed based on the loci A, B, and DRB1.

### Cellular reprogramming and iPS generation

#### Ethical Approval

This study was approved by the Institutional Review Board of King Abdullah International Medical Research Center (KAIMRC) (Protocol# RJ20/134/J). Initial donor recruitment was done by the Saudi Stem Cell Donor Registry staff. Personal interview was conducted, and informed consents were obtained by the research team.

### PBMCs isolation and enrichment of erythroid progenitors

Peripheral blood was collected from Saudi patients into an EDTA-containing blood collection tube and treated with RosetteSep^TM^ Human Progenitor Cell Basic Pre-Enrichment antibody cocktail according to the manufacturer’s instructions (Stem Cell Technologies Catalog#15226). After PBMCs separation and isolation, 1 million cells were cultured for 8 days in StemSpan^TM^ SFEM II medium (Stem Cell Technologies Catalog #09605) supplemented with 1X StemSpan^TM^ Erythroid Expansion Supplement (Stem Cell Technologies Catalog #02692).

### Transfection of Erythroid Progenitor Cells

Expanded erythroid cells were reprogrammed with Episomal iPSC Reprogramming Kit (Thermofisher Catalog#A15960). Around 3 x 105 cells were electroporated with 1 μg of each episomal vector using Neon Transfection System (Thermofisher). The emerging ESC-like colonies were manually picked and transferred into 96-well plates coated with rhLaminin-521 (Thermofisher Catalog#A29248) in mTeSR™ Plus medium (Stem Cell Technologies Catalog #100-0276). iPSCs were dissociated ReLeSR (Thermofisher Catalog # 100-0484) using 1:10-1:30 splitting ratio and incubated at 37°C, 5% CO2, 20% O2 incubator.

### Molecular characterization of pluripotency and genomic integrity

#### Immunocytochemistry

Cells were fixed in 4% (w/v) paraformaldehyde for 15 min, permeabilized in PBS containing 0.1% (v/v) Triton X-100 for 10 min, and subsequently blocked in PBS containing 1% Gelatin for 45 min. Cells were incubated with primary antibodies overnight at 4 °C and probed with the appropriate secondary antibodies for 1 hour at room temperature (ThermoFisher Scientific). Primary and secondary antibodies were resuspended in 0.2% Gelatin in PBS. The nuclei were counterstained with 1 μg /mL DAPI nuclear staining (Thermo Fisher Scientific).

### Quantitative Reverse Transcription PCR (RT-qPCR)

Total RNA was extracted using RNeasy Kit (Qiagen Catalog# 74104) and reverse transcribed using the High-Capacity cDNA Reverse Transcription Kit (Applied Biosystems^TM^ Catalog# 4374966). The RT-qPCR assay was carried out using FastStart SYBR Green Master Mix (ROCHE) as described previously (Alsayegh et al., 2018).

### In-vitro Differentiation

The generated iPSCs were differentiated into three germ layers using the STEMdiff^TM^ Trilineage Differentiation Kit (Stem Cell Technologies Catalog #05230).

### Flow Cytometry Analyses

Cells were stained with OCT4, NANOG, SOX2, and cTnI antibodies diluted in 2% FBS in PBS for 30 mins on ice in the dark with occasional vortexing. It was then washed with PBS and analyzed on BD FACS ARIA cell sorter. FITC-positive cells were measured in stained vs unstained cells.

### Karyotyping

For G banding karyotyping, iPSC lines were treated with 0.3 μg/mL KaryoMAX^TM^ Colcemid^TM^ (1 μg) for 15 min, dissociated by TrypLE, and incubated in hypotonic solution (75 mM potassium chloride) at 37 °C for 20 min. iPSCs were then fixed in methanol/glacial acetic acid 3:1 and stored at 4 °C. At least 50 metaphases were karyotyped at the department of pathology and laboratory medicine (Ministry of the National Guard – Health Affairs).

### Neural Progenitor Cells (NPCs) differentiation

The generation of central nervous system (CNS)-type neural progenitor cells (NPCs) from HLA-iPSCs was performed according to Monolayer Culture Protocol (STEMdiff^TM^ SMADi Neural Induction Kit Catalog #08581).

### Cardiomyocyte differentiation

The differentiation of hESCs toward beating cardiomyocyte was performed following STEMdiff^TM^ Ventricular Cardiomyocyte Differentiation Kit (Stem Cell Technologies Catalog #05010) in accordance with the manufacturer instructions. In brief, iPSCs were detached using gentle cell dissociation reagent and seeded at 1.2 x 10^6^ cells/well on Matrigel-coated 12 well plates in presence of mTeSR^TM^ Plus medium and 10 μM Y-27632. Subsequently, the differentiation was initiated by replacing culture medium with Cardiomyocyte Differentiation Medium A for 48 hrs. at 37 ◦C, 5%CO2. Cardiomyocyte Differentiation Medium B was added for another 48 hrs. Then, Cardiomyocyte Differentiation Medium C was replaced on day 4 and 6. We perform a full-medium change with Cardiomyocyte Maintenance Medium every other day up to 20 days.

### Episomal Plasmids Screening

DNA was extracted using AllPrep DNA/RNA/ Mini Kit (Qiagen Catalog# 80204). PCR was performed using EBNA-1 primers that detect all five episomal plasmids (expected size 666 bp) according to manufacture guidelines (Thermo Fisher Scientific Catalog # A15960).

### Mycoplasma Detection

Mycoplasma contamination was assessed using LookOut^®^ Mycoplasma qPCR Detection (SIGMA) (Supplementary Figure 1C).

### Statistical Analysis

RT-qPCR data are represented as mean ± standard deviation (SD). Statistical significance was determined in Student’s *t*-test (unpaired; two-tailed). A Bonferroni correction was applied to the p-value from multiple comparisons. *p < 0.05.

### Short Tandem Repeats (STR) Identity Assay

Extracted genomic DNA from HLA-iPSCs and PBMCs was analyzed for 24 polymorphic STR markers using GenePrint 24 system (Promega, Madison, USA) following manufacturer’s protocol and were amplified using PCR and followed by ABI capillary electrophoresis. In this analysis, a 24 autosomal STR were analysed D8S1179, D21S11, D7S820, CSF1PO, D3S1358, THO1, D13S317, D16S539, D2S1338, D19S433, vWA, TPOX, D18S51, D5S818, D1S1656, D2S441, D10S1248, Penta E, Penta D, DYS391, D12S391, D22S1045 FGA and Amelogenin using ABI 3130/3500 Genetic Analyzer.

### Whole Genome Sequencing (WGS)

The Nextera library prep kit (Illumina) was used to prepare libraries for WGS resequencing on the Novaseq 6000 sequencer (Illumina). The short-read sequences obtained from a blood sample as control, as well as the two cell lines iPSC#1 and iPSC#2 were assessed with FastQC (Andrews, n.d.). Adapter and low quality regions were trimmed with Trimmomatic v0.33 (Bolger et al., 2014) using parameters: 2:30:10 LEADING:3 TRAILING:3 SLIDINGWINDOW:4:20 MINLEN:40, leaving 280, 262, and 245 million reads for the blood, iPSC#1 and iPSC#2 dataset respectively. Trimmed reads were mapped to the human reference genome assembly UCSC hg38 analysis set using BWA mem (Li & Durbin, 2010), which yielded a median coverage between 23X and 26X. Duplicates were marked and read groups added with Picard tools (Broad Institute, 2018). The donor sample as well as both generated cell lines were HLA genotyped using xHLA (https://www.pnas.org/doi/10.1073/pnas.1707945114). Single nucleotide polymorphisms were jointly called in all samples with GATK HaplotypeCaller (McKenna et al., 2010) following GATK best practices recommendations as well as with GATK Mutect2. In case of Mutect, mapped reads from HLA-iPSC#1 and HLA-iPSC#2 were run separately as treatment while the blood sample was provided as normal reference. Single nucleotide polymorphism calls were then filtered requiring a minimal allele frequency of 20%. The obtained polymorphisms were then annotated using the Ensembl Variant Effect Predictor VEP (McLaren, Genome Biol. 2016). Detected SNVs were tested for overlap with genes listed in the Catalogie of Somatic Mutations In Cancer (COSMIC) Cancer Gene Census (https://cancer.sanger.ac.uk/census) and the Shibata cancer gene panel (https://www.pmda.go.jp/files/000152599.pdf). Variants which were predicted to have a high impact on the aforementioned gene set were manually examined.

Structural variants were called with Delly v1.1.6 and Manta v1.6 in subtractive mode specifying the blood sample as control and both cell lines separately as treatments. Structural variants passing the quality filter for each caller and being classified as “precise” were retained. Variants calls for each cell line from Delly and Manta were then compared using SURVIVOR (Jeffares et al. 2017, Nat. Comm.) and only overlapping calls were retained for further analysis, allowing for at most 500bp distance between break points. This procedure revealed no structural variants in HLA-iPSC#2 and one tandem duplication HLA-iPSC#1.

## Notes

### Competing Interest Statement

The authors have declared no competing interest.

