## Supplementary Data for "HLA-Based Banking of Human Induced Pluripotent Stem Cells in Saudi Arabia"

### Slide 1
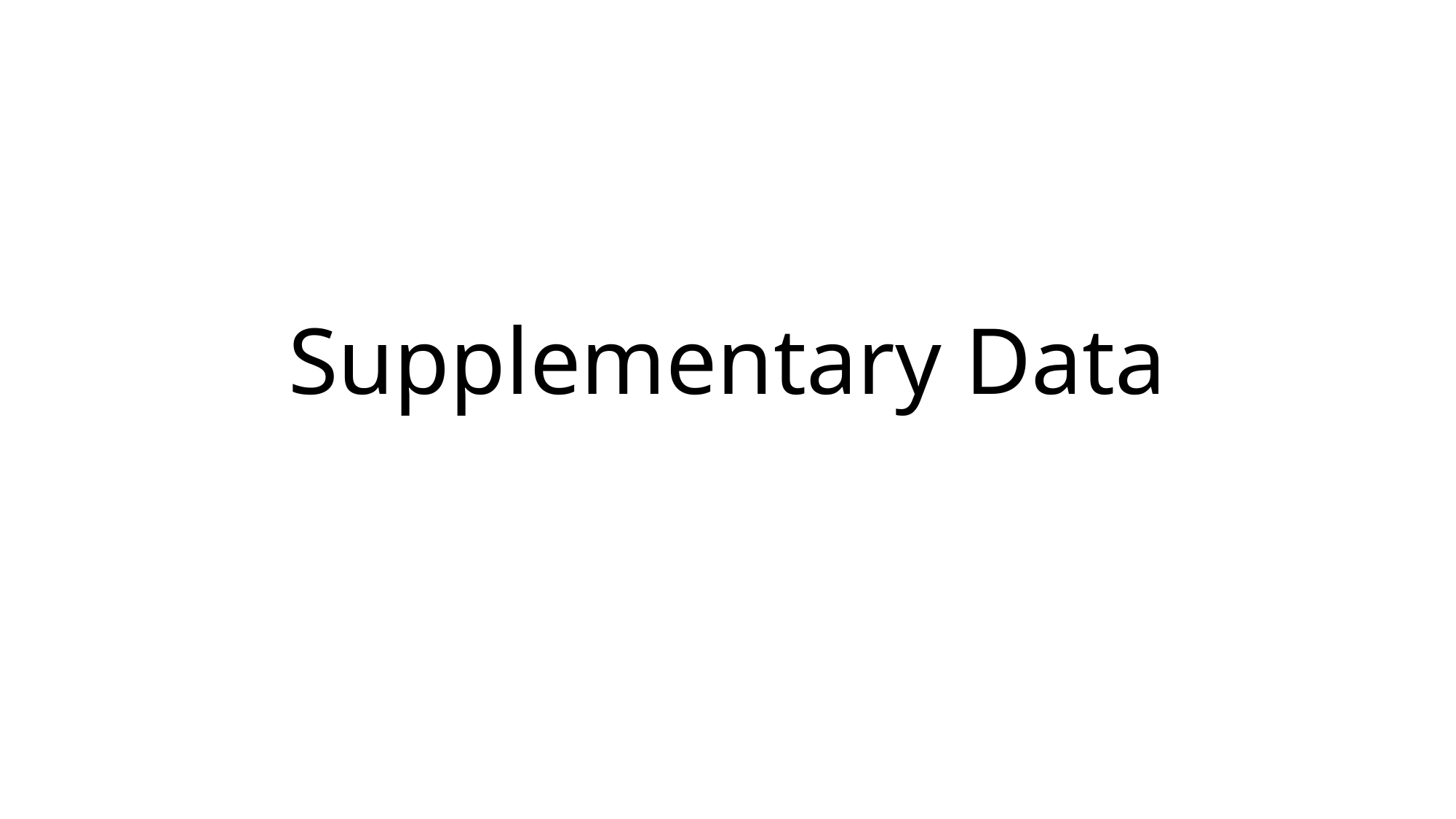

#### Supplementary Data

### Slide 2
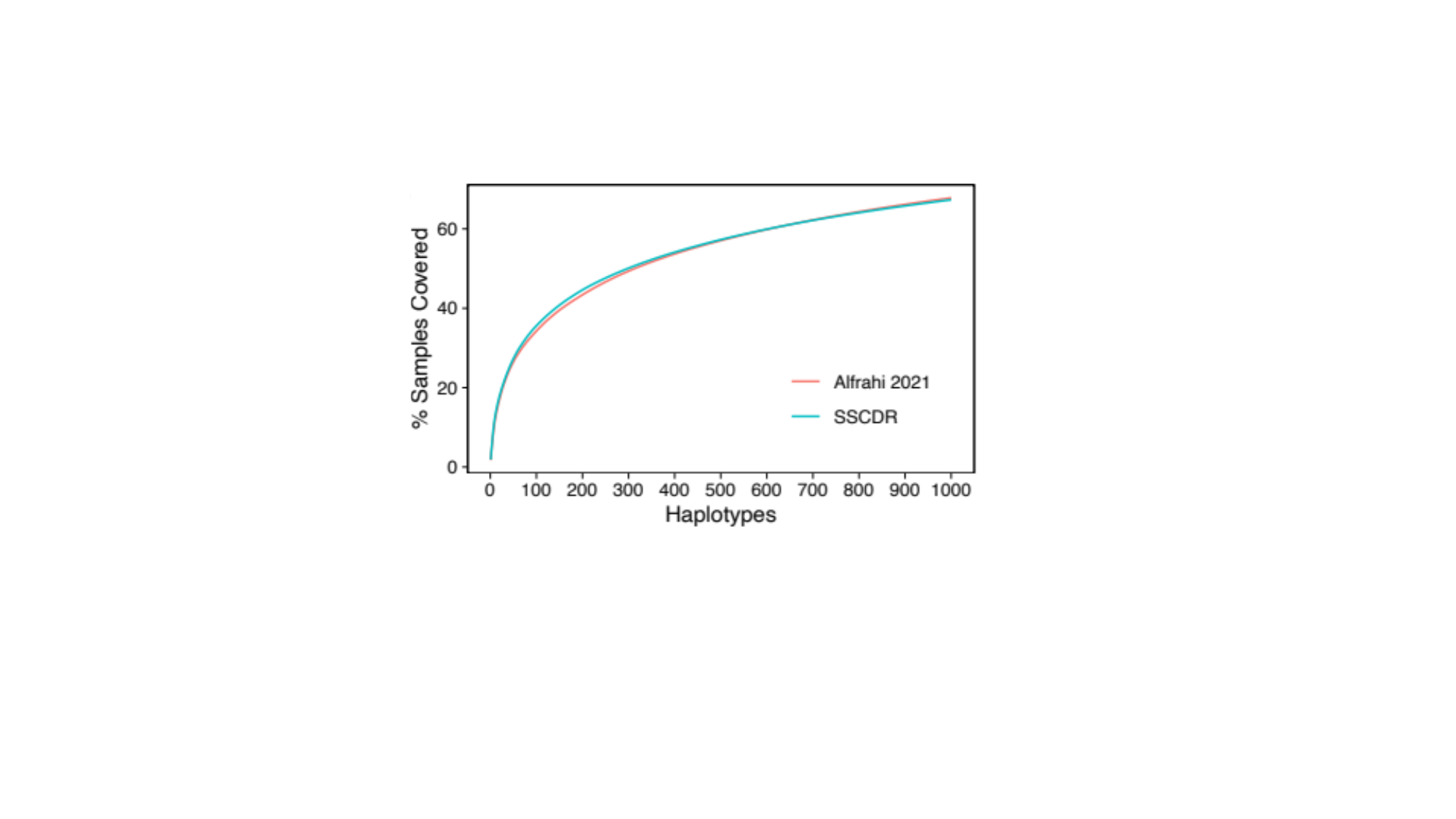

### Slide 3
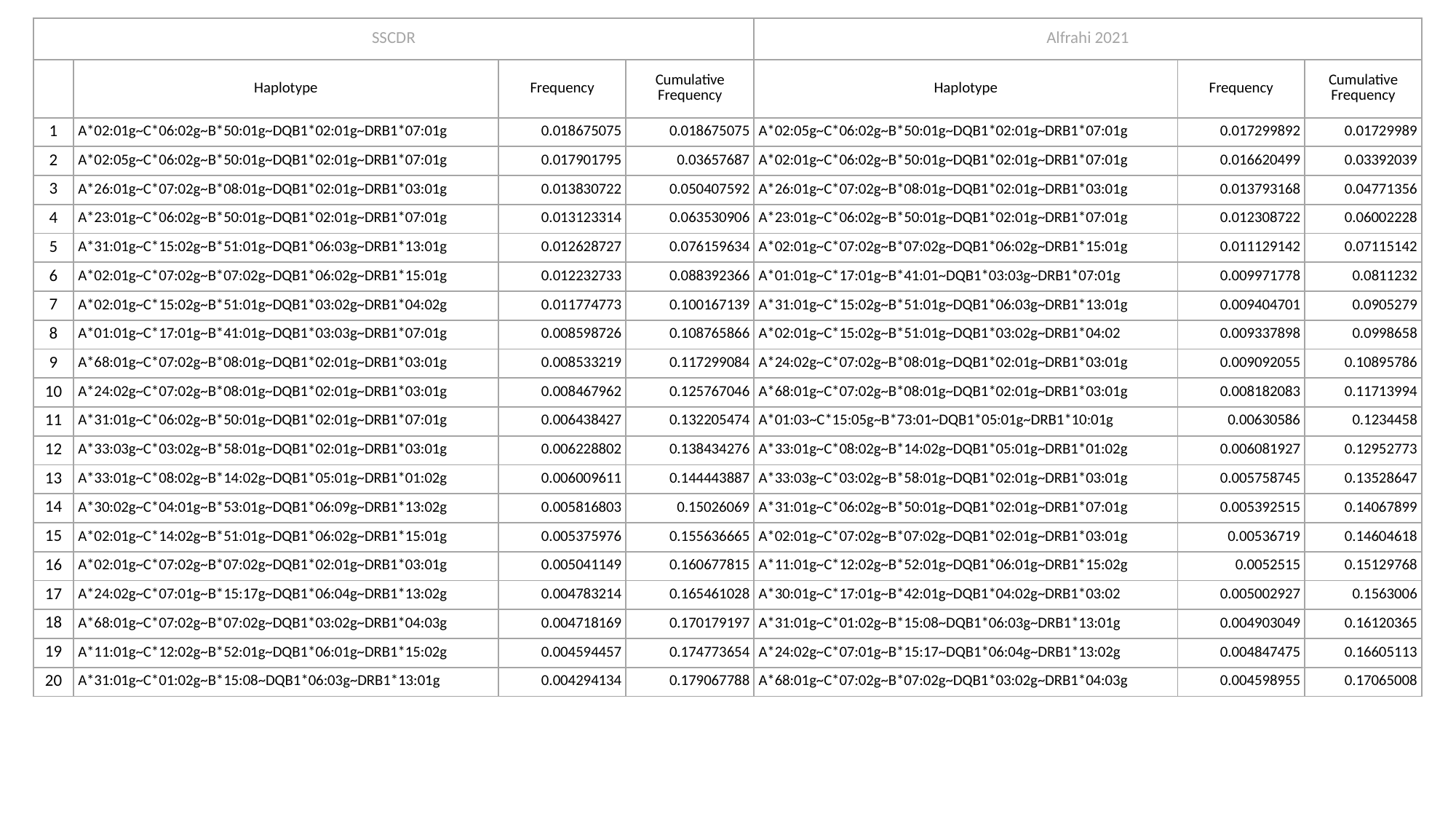

| SSCDR | | | | Alfrahi 2021 | | |
| --- | --- | --- | --- | --- | --- | --- |
| | Haplotype | Frequency | Cumulative Frequency | Haplotype | Frequency | Cumulative Frequency |
| 1 | A\*02:01g~C\*06:02g~B\*50:01g~DQB1\*02:01g~DRB1\*07:01g | 0.018675075 | 0.018675075 | A\*02:05g~C\*06:02g~B\*50:01g~DQB1\*02:01g~DRB1\*07:01g | 0.017299892 | 0.01729989 |
| 2 | A\*02:05g~C\*06:02g~B\*50:01g~DQB1\*02:01g~DRB1\*07:01g | 0.017901795 | 0.03657687 | A\*02:01g~C\*06:02g~B\*50:01g~DQB1\*02:01g~DRB1\*07:01g | 0.016620499 | 0.03392039 |
| 3 | A\*26:01g~C\*07:02g~B\*08:01g~DQB1\*02:01g~DRB1\*03:01g | 0.013830722 | 0.050407592 | A\*26:01g~C\*07:02g~B\*08:01g~DQB1\*02:01g~DRB1\*03:01g | 0.013793168 | 0.04771356 |
| 4 | A\*23:01g~C\*06:02g~B\*50:01g~DQB1\*02:01g~DRB1\*07:01g | 0.013123314 | 0.063530906 | A\*23:01g~C\*06:02g~B\*50:01g~DQB1\*02:01g~DRB1\*07:01g | 0.012308722 | 0.06002228 |
| 5 | A\*31:01g~C\*15:02g~B\*51:01g~DQB1\*06:03g~DRB1\*13:01g | 0.012628727 | 0.076159634 | A\*02:01g~C\*07:02g~B\*07:02g~DQB1\*06:02g~DRB1\*15:01g | 0.011129142 | 0.07115142 |
| 6 | A\*02:01g~C\*07:02g~B\*07:02g~DQB1\*06:02g~DRB1\*15:01g | 0.012232733 | 0.088392366 | A\*01:01g~C\*17:01g~B\*41:01~DQB1\*03:03g~DRB1\*07:01g | 0.009971778 | 0.0811232 |
| 7 | A\*02:01g~C\*15:02g~B\*51:01g~DQB1\*03:02g~DRB1\*04:02g | 0.011774773 | 0.100167139 | A\*31:01g~C\*15:02g~B\*51:01g~DQB1\*06:03g~DRB1\*13:01g | 0.009404701 | 0.0905279 |
| 8 | A\*01:01g~C\*17:01g~B\*41:01g~DQB1\*03:03g~DRB1\*07:01g | 0.008598726 | 0.108765866 | A\*02:01g~C\*15:02g~B\*51:01g~DQB1\*03:02g~DRB1\*04:02 | 0.009337898 | 0.0998658 |
| 9 | A\*68:01g~C\*07:02g~B\*08:01g~DQB1\*02:01g~DRB1\*03:01g | 0.008533219 | 0.117299084 | A\*24:02g~C\*07:02g~B\*08:01g~DQB1\*02:01g~DRB1\*03:01g | 0.009092055 | 0.10895786 |
| 10 | A\*24:02g~C\*07:02g~B\*08:01g~DQB1\*02:01g~DRB1\*03:01g | 0.008467962 | 0.125767046 | A\*68:01g~C\*07:02g~B\*08:01g~DQB1\*02:01g~DRB1\*03:01g | 0.008182083 | 0.11713994 |
| 11 | A\*31:01g~C\*06:02g~B\*50:01g~DQB1\*02:01g~DRB1\*07:01g | 0.006438427 | 0.132205474 | A\*01:03~C\*15:05g~B\*73:01~DQB1\*05:01g~DRB1\*10:01g | 0.00630586 | 0.1234458 |
| 12 | A\*33:03g~C\*03:02g~B\*58:01g~DQB1\*02:01g~DRB1\*03:01g | 0.006228802 | 0.138434276 | A\*33:01g~C\*08:02g~B\*14:02g~DQB1\*05:01g~DRB1\*01:02g | 0.006081927 | 0.12952773 |
| 13 | A\*33:01g~C\*08:02g~B\*14:02g~DQB1\*05:01g~DRB1\*01:02g | 0.006009611 | 0.144443887 | A\*33:03g~C\*03:02g~B\*58:01g~DQB1\*02:01g~DRB1\*03:01g | 0.005758745 | 0.13528647 |
| 14 | A\*30:02g~C\*04:01g~B\*53:01g~DQB1\*06:09g~DRB1\*13:02g | 0.005816803 | 0.15026069 | A\*31:01g~C\*06:02g~B\*50:01g~DQB1\*02:01g~DRB1\*07:01g | 0.005392515 | 0.14067899 |
| 15 | A\*02:01g~C\*14:02g~B\*51:01g~DQB1\*06:02g~DRB1\*15:01g | 0.005375976 | 0.155636665 | A\*02:01g~C\*07:02g~B\*07:02g~DQB1\*02:01g~DRB1\*03:01g | 0.00536719 | 0.14604618 |
| 16 | A\*02:01g~C\*07:02g~B\*07:02g~DQB1\*02:01g~DRB1\*03:01g | 0.005041149 | 0.160677815 | A\*11:01g~C\*12:02g~B\*52:01g~DQB1\*06:01g~DRB1\*15:02g | 0.0052515 | 0.15129768 |
| 17 | A\*24:02g~C\*07:01g~B\*15:17g~DQB1\*06:04g~DRB1\*13:02g | 0.004783214 | 0.165461028 | A\*30:01g~C\*17:01g~B\*42:01g~DQB1\*04:02g~DRB1\*03:02 | 0.005002927 | 0.1563006 |
| 18 | A\*68:01g~C\*07:02g~B\*07:02g~DQB1\*03:02g~DRB1\*04:03g | 0.004718169 | 0.170179197 | A\*31:01g~C\*01:02g~B\*15:08~DQB1\*06:03g~DRB1\*13:01g | 0.004903049 | 0.16120365 |
| 19 | A\*11:01g~C\*12:02g~B\*52:01g~DQB1\*06:01g~DRB1\*15:02g | 0.004594457 | 0.174773654 | A\*24:02g~C\*07:01g~B\*15:17~DQB1\*06:04g~DRB1\*13:02g | 0.004847475 | 0.16605113 |
| 20 | A\*31:01g~C\*01:02g~B\*15:08~DQB1\*06:03g~DRB1\*13:01g | 0.004294134 | 0.179067788 | A\*68:01g~C\*07:02g~B\*07:02g~DQB1\*03:02g~DRB1\*04:03g | 0.004598955 | 0.17065008 |

### Slide 4
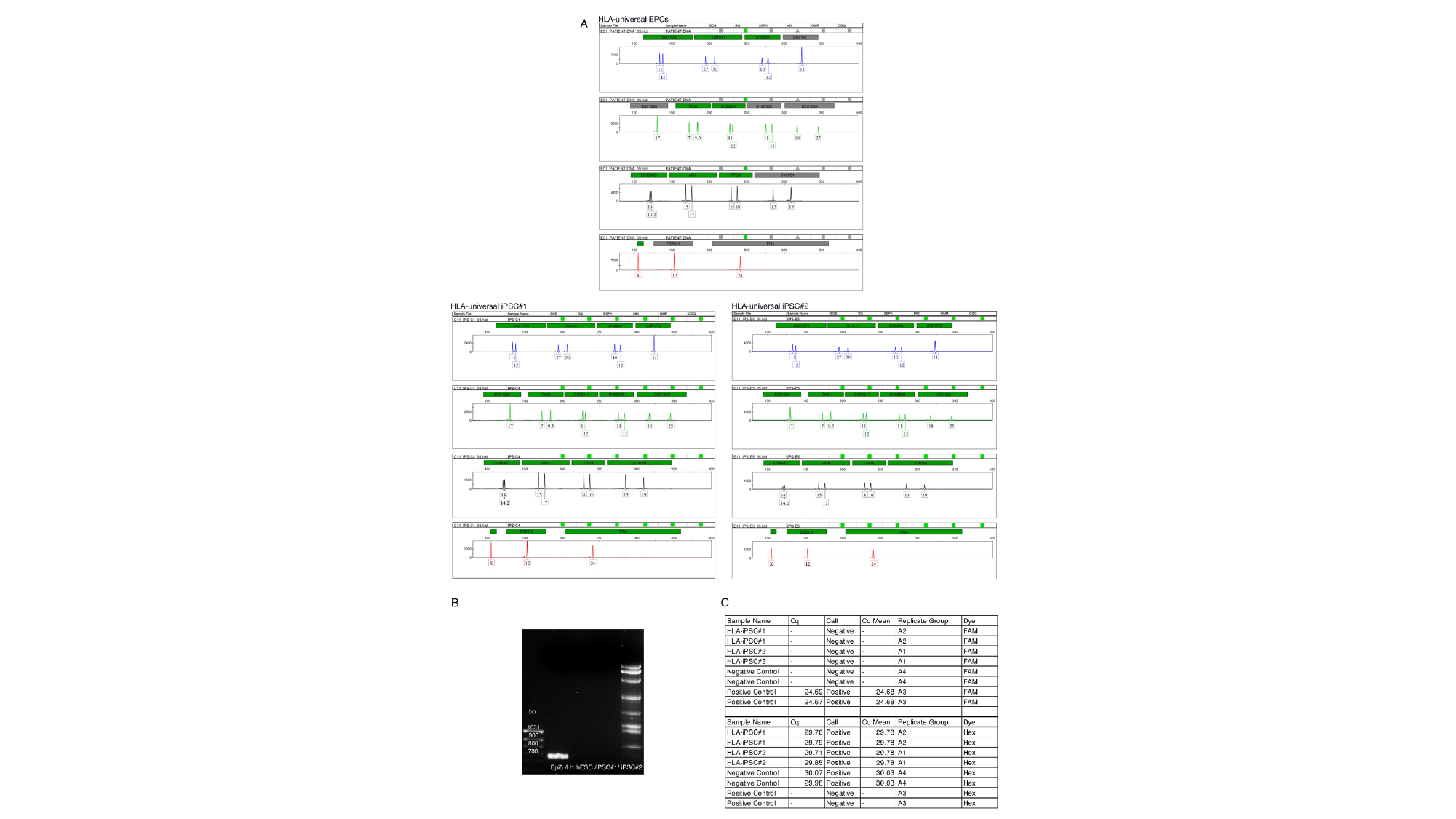

### Slide 5
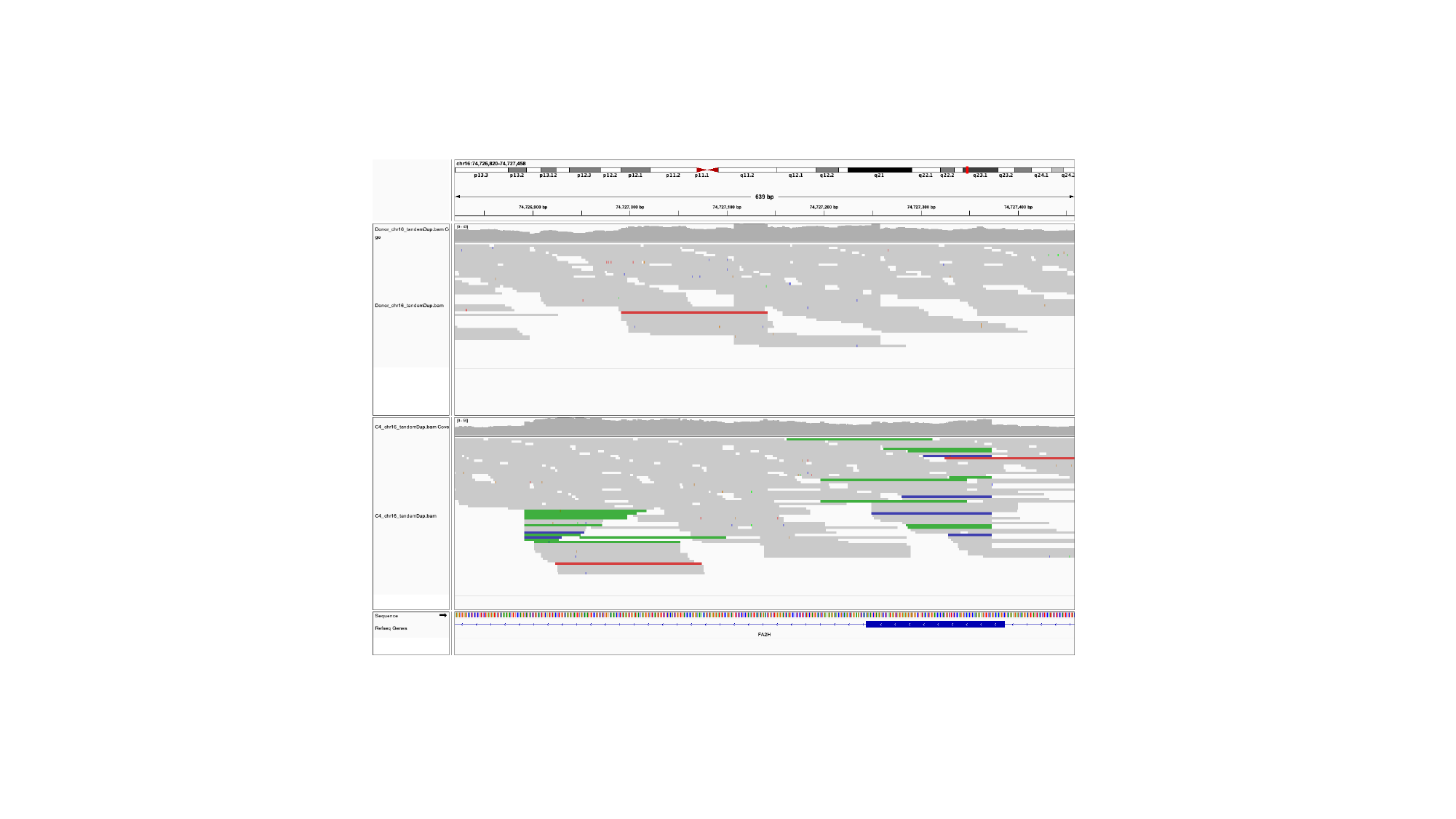

### Slide 6
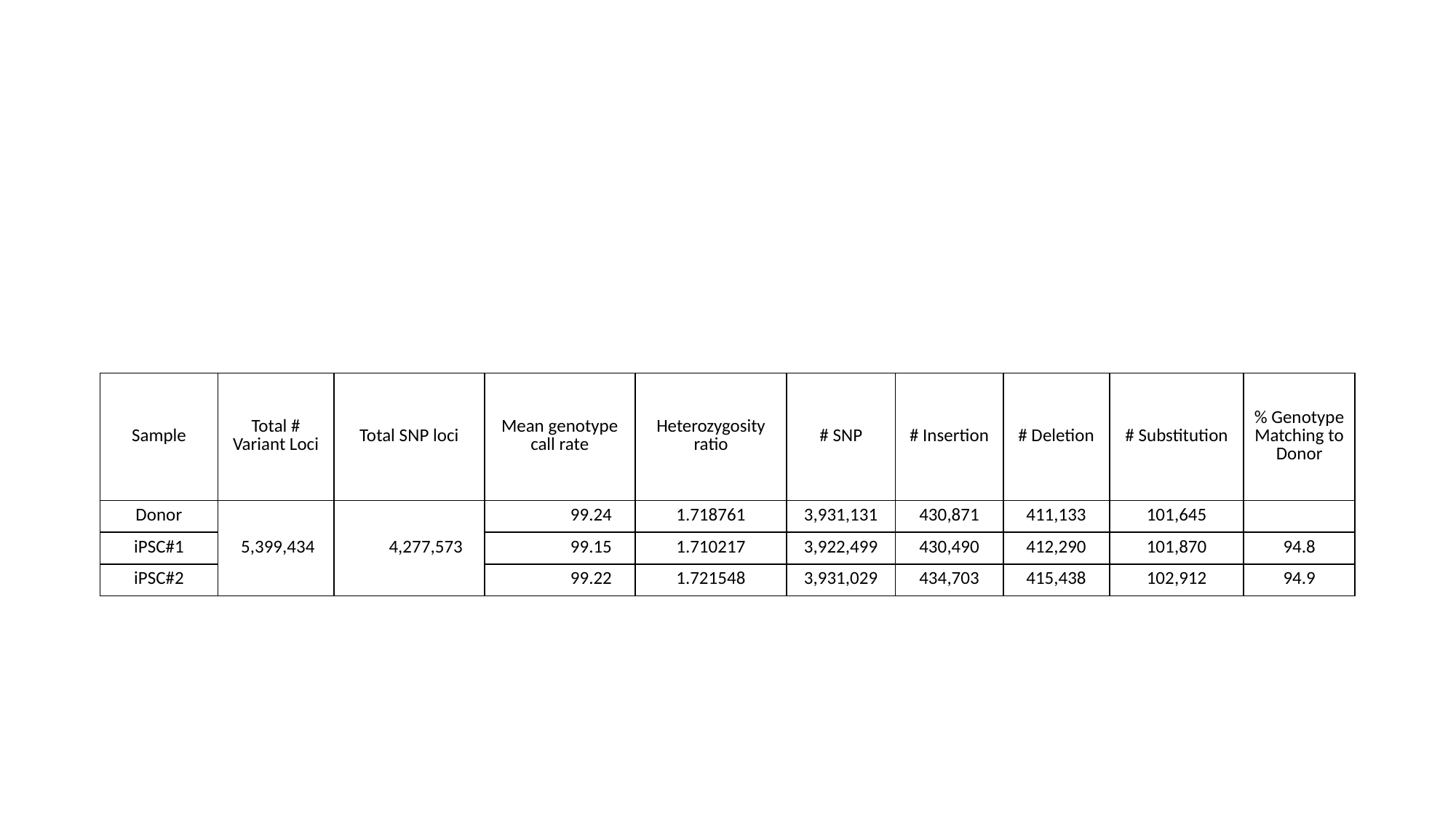

| Sample | Total # Variant Loci | Total SNP loci | Mean genotype call rate | Heterozygosity ratio | # SNP | # Insertion | # Deletion | # Substitution | % Genotype Matching to Donor |
| --- | --- | --- | --- | --- | --- | --- | --- | --- | --- |
| Donor | 5,399,434 | 4,277,573 | 99.24 | 1.718761 | 3,931,131 | 430,871 | 411,133 | 101,645 | |
| iPSC#1 | | | 99.15 | 1.710217 | 3,922,499 | 430,490 | 412,290 | 101,870 | 94.8 |
| iPSC#2 | | | 99.22 | 1.721548 | 3,931,029 | 434,703 | 415,438 | 102,912 | 94.9 |

### Slide 7
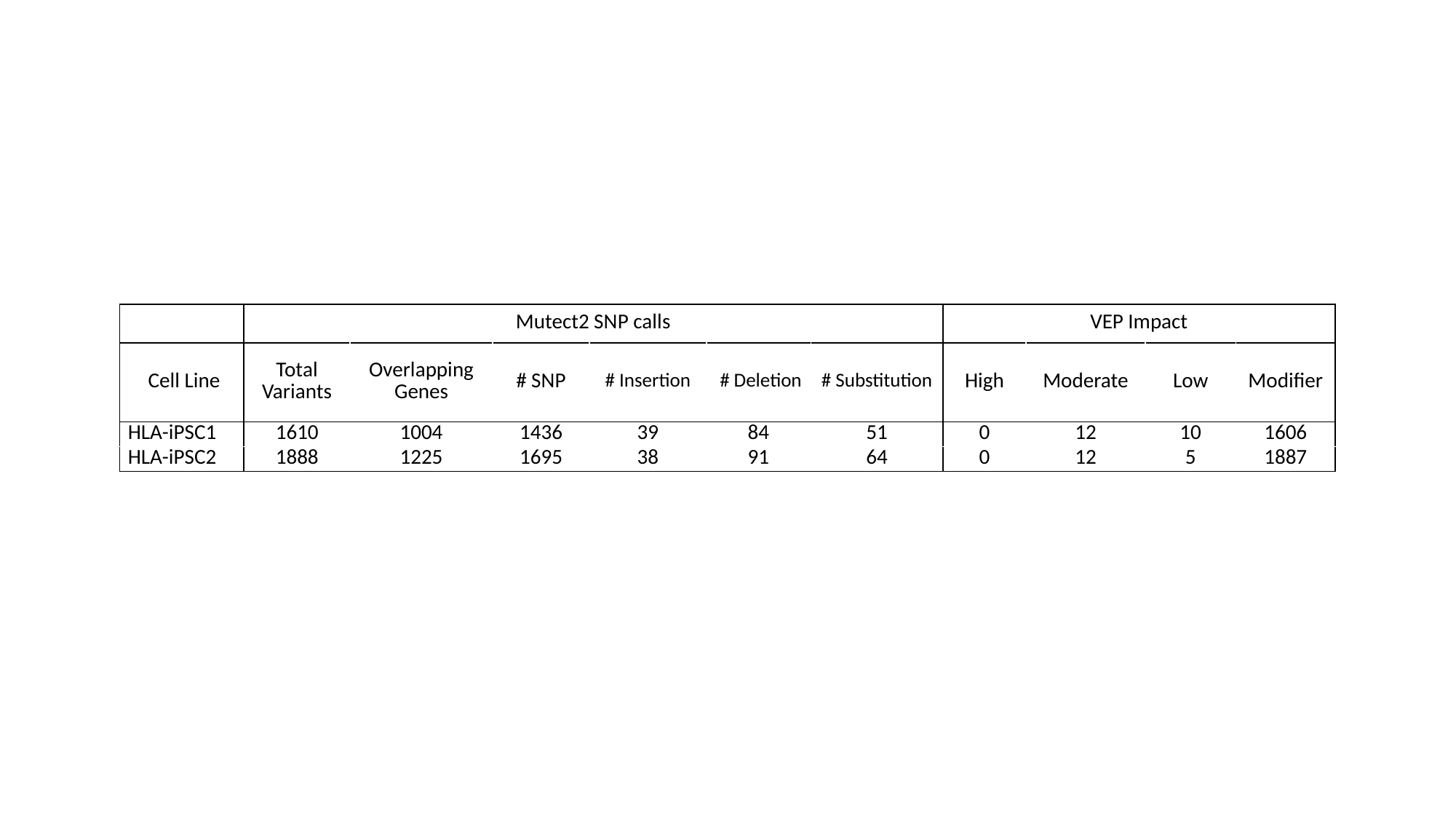

#
| | Mutect2 SNP calls | | | | | | VEP Impact | | | |
| --- | --- | --- | --- | --- | --- | --- | --- | --- | --- | --- |
| Cell Line | Total Variants | Overlapping Genes | # SNP | # Insertion | # Deletion | # Substitution | High | Moderate | Low | Modifier |
| HLA-iPSC1 | 1610 | 1004 | 1436 | 39 | 84 | 51 | 0 | 12 | 10 | 1606 |
| HLA-iPSC2 | 1888 | 1225 | 1695 | 38 | 91 | 64 | 0 | 12 | 5 | 1887 |
